# Effect of leaf temperature on estimating physiological traits of wheat leaves from hyperspectral reflectance

**DOI:** 10.1101/2020.05.21.109652

**Authors:** Hammad A Khan, Yukiko Nakamura, Robert T Furbank, John R Evans

**Author notes:** CSIRO Agriculture & Food, Waite Campus, Glen Osmond, SA 5064, Australia.

## Abstract

A growing number of leaf traits can be predicted from hyperspectral reflectance data. These include structural and compositional traits, such as leaf mass per area, nitrogen and chlorophyll content, but also physiological traits such a Rubisco carboxylation activity, electron transport rate and respiration rate. Since physiological traits vary with leaf temperature, how does this impact on predictions made from reflectance measurements? We investigated this with two wheat varieties, by repeatedly measuring each leaf through a sequence of temperatures imposed by varying the air temperature in a growth room. The function predicting Rubisco capacity normalised to 25 °C predicted the same value, regardless of leaf temperatures ranging from 20 to 35°C. Leaf temperature affected none of the predicted traits: V_cmax25_, J, chlorophyll content, LMA, N content per unit leaf area or V_cmax25_/N. However, as others have derived models to predict Rubisco activity that includes variation associated with leaf temperature, we discuss whether these functions may include a temperature signal within the reflectance spectra.

## Introduction

Plant breeders continually strive to improve crop yield. For cereals, there has been a recognition that future increases could benefit from improving photosynthesis (Parry *et al*., 2011; Reynolds *et al*., 2009). Crop growth is not simply related to a measurement of photosynthetic rate of a particular leaf under one condition. Instead, photosynthesis integrated over a day with contributions from all the leaves in the canopy drives crop growth. Subsequent conversion into biomass and the partitioning into harvested grains determines yield. All these processes combined pose a major challenge on how to meaningfully measure photosynthesis with the goal of improving yield. However, there are a few examples that have compared historical sequences of cultivars and observed correlations between leaf photosynthetic rate and wheat yield (Beche *et al*., 2014; Fischer *et al*., 1998; Gaju *et al*., 2016; Yao *et al*., 2019). It has also been found that radiation use efficiency (above ground biomass produced per unit of intercepted photosynthetically active radiation) has been increasing over time with changing wheat varieties in both the UK (Shearman *et al*., 2005) and Australia (Sadras *et al*., 2012). Interestingly, both studies found the same rate of increase (0.012 g MJ^−1^ y^−1^).

It is possible to survey photosynthetic properties between wheat genotypes (Driever *et al*., 2014; Silva-Pérez *et al*., 2019), but detailed phenotyping is time-consuming which limits the number of genotypes that can be sampled. A promising alternative is to predict photosynthetic traits from leaf reflectance spectra. Serbin et al. (2012) derived models predicting nitrogen concentration, leaf dry mass per unit area, maximum Rubisco carboxylase activity (V_cmax_) and photosynthetic electron transport rate (J) from hyperspectral reflectance measured on leaves of *Populus tremuloides* and *P. deltoides*. Leaf temperature varied between 20 and 30 °C, depending on the glasshouse regime, which strongly influenced V_cmax_. A single model was presented that applied to both species, regardless of leaf temperature. It was argued that this implied that the derived V_cmax_ was not being predicted indirectly from another trait such as nitrogen (Serbin *et al*., 2012). The hyperspectral reflectance approach was also successfully used to predict V_cmax_ for *Glycine max* measured between 26 and 34°C (Ainsworth *et al*., 2014) and *Nicotiana tabacum* (Meacham-Hensold *et al*., 2019) and *Zea mays* (Yendrek *et al*., 2017) measured at various temperatures in the field.

In order to be able to make useful comparisons of Rubisco capacity between plants which may differ in their leaf temperature during sampling, one needs to know both V_cmax_ and leaf temperature. Alternatively, one could use the temperature responses of the Rubisco enzyme kinetic parameters (Bernacchi *et al*., 2003; Silva-Perez *et al*., 2017) to convert gas exchange estimates of V_cmax_ to a common temperature, e.g. 25°C (V_cmax25_), which are then used to build a model from reflectance data. This has been done for a group of 37 broadleaf tree species (Dechant *et al*., 2017), wheat (Silva-Perez *et al*., 2018) and 21 tropical tree species from Panama and Brazil (Wu *et al*., 2019). Heckmann et al. (2017) also presented predictions of V_cmax_ for *Brassica, Moricandia* and Z. *mays* from reflectance spectra, but measured only at 25°C. While Dechant et al. (2017) normalised their gas exchange to 25°C, the reflectance spectra were collected at prevailing leaf temperatures. It is not known whether the prediction of V_cmax25_ from leaf reflectance is insensitive to the temperature of the leaf during the reflectance measurement. We, therefore, set out to specifically assess whether leaf trait predictions from hyperspectral reflectance varied with leaf temperature by repeatedly measuring the same leaf sequentially through a range of temperatures in two wheat cultivars. We hypothesized that leaf temperature would not affect predicted values of leaf traits obtained using leaf hyperspectral reflectance.

## Materials and Methods

### Plant material and growth conditions

Expt 1 Two spring wheat genotypes *(Triticum aestivum* Kukri and Seri) were grown in a naturally lit greenhouse (day/night temperatures set at 25/15 °C) at the Australian National University in Canberra during Sep-Nov 2018. Three seeds were sown in well-drained 3.5-litre pots filled with commercial potting mix, containing basal fertilizer Osmocote (Scotts). Pots were laid out according to randomized block design with six replicates and blocks representing the replications. After emergence, seedlings were thinned down to one plant per pot. Plants were watered daily until the end of the experiment.

Temperature treatment was given in a controlled environmental chamber, with day/night temperatures set at 25/15 °C and irradiance set to 500 μmol photons m^−2^ s^−1^. All the measurements were made seven days after anthesis. Plants were moved to the chamber one day before the actual measurements so that plants could acclimatize to the chamber’s environmental conditions. The next day, measurements were made at a chamber temperature of 15, 25, 35 and 15 °C, in the described sequence. After achieving the desired chamber temperature, plants were acclimatized at least 1 hr before the measurements were made.

Expt 2 Two spring wheat genotypes (Kukri and Seri) and one triticale (Hawkeye) were grown in a greenhouse with temperature set to 20/15 °C (day/night). Seeds were sown on multiple days in March 2018 and each genotype was sown separately in shallow tray with raising mix. After germination, seedlings were transplanted into 5L pots filled commercial potting mix, containing basal fertilizer Osmocote (Scotts). Five-six weeks after sowing, half of the plants were transferred to an adjacent greenhouse room set at 32/20°C, where they grew for one-two more weeks before gas exchange measurements were made.

### Hyperspectral reflectance measurements (Expt 1)

Hyperspectral reflectance spectra were measured with a FieldSpec^®^4 (Analytical Spectral Devices, Boulder, CO, USA) full range spectroradiometer (350–2500 nm) attached to a leaf clip (Analytical Spectral Devices, Boulder, CO, USA) with a fibre optic cable. Leaf clip had an internal calibrated light source and two external panels i.e. a white panel to calibrate the instrument and a black panel for taking measurements. A mask containing a black circular gasket was also attached to leaf clip, which was used to reduce the leaf-clip aperture to an oval area (1.15 x 1.4 cm = 1.264 cm^2^) suitable for a wheat leaf (Silva Perez ref). For each temperature, one reflectance measurement was made at the same place of the flag leaf of each plant by putting the leaf vertically to the leaf probe as explained elsewhere (Silva-Perez *et al*., 2018).

Leaf reflectance spectra were processed according to Silva-Perez *et al*. (2018). A ‘jump’ correction associated with a change in the detectors at 1000 and 1800 nm was applied before the traits were predicted.

### MultispeQ. measurements (Expt 1)

Linear electron transport (LET) and relative chlorophyll content (SPAD units) measurements were carried out using a handheld MultispeQ (Beta) device linked to the PhotosynQ platform (www.photosynq.org) (Kuhlgert *et al*., 2016). Relative chlorophyll content (SPAD units) was estimated by measuring the transmittance of red (650 nm) and infrared (940 nm) light. LET was estimated from the measurements of quantum yield of photosystem II (■11) via pulse-amplitude modulation (PAM) fluorometry at photosynthetically active radiation (PAR) of 1000 μmol photons m^−2^ s^−1^ (Kuhlgert *et al*., 2016).

### Gas-exchange measurements (Expt 2)

Gas exchange was measured on the most recently fully expanded leaves with a LI-6400XT Portable Photosynthesis system (LI-COR Biosciences Inc., Lincoln, NE, USA) on plants placed inside a controlled environment cabinet (Thermoline Science Model-TRIL/SL). The air flow rate was 500 μmol s^−1^ with a PPFD of 1800 μmol m^−2^ s^−1^ supplied by the LED light. Gas exchange was measured at leaf temperatures of 15, 25 and 35°C. At each temperature, CO_2_ response curves were measured in 21% O_2_ using inlet CO_2_ concentrations of 400, 50, 100, 150, 250, 400, 600, 800, 1000, 400 μmol mol^1^. Subsequently, the air was changed to 2% O_2_ with a CO_2_ concentration in the leaf chamber of 380 μmol mol^−1^, the flow reduced to 200 μmol s^−1^ and measurement continued for 60 minutes with concurrent sampling for carbon isotope discrimination to determine mesophyll conductance (Evans and von Caemmerer, 2013). Maximum Rubisco carboxylase activity (*V_cmax_*) was calculated from CO_2_ response curves using kinetic constants derived from wheat (Silva-Pérez *et al*., 2017).

### Statistical analysis

Data were subjected to analysis of variance using various packages in R (R coreR, 2013). Means were compared for significant differences using Tukey’s multiple comparison tests at 5% probability level.

## Results

The consequence of using leaf reflectance spectra collected from leaves with varying temperatures to predict leaf traits was investigated with two wheat varieties.

The predicted value of V_cmax25_ was not affected by the leaf temperature when reflectance spectra were measured, for either cultivar (Fig. 1A). Upon returning the growth cabinet to 15°C, the V_cmax25_ values were not significantly different from the initial values. By contrast, V_cmax_ values derived from gas exchange increased fourfold between 15 and 35°C (Fig. 1B). Values for wheat grown under cool or hot conditions showed a difference at 35°C, with the cool grown plants falling further below the theoretical line (consistent with an E_a_ of 63kJ mol^−1^) than plants from the hot treatment. The values for plants from the hot treatment superimpose the previously published data from Silva-Perez et al. (2017).

**Fig. 1.**
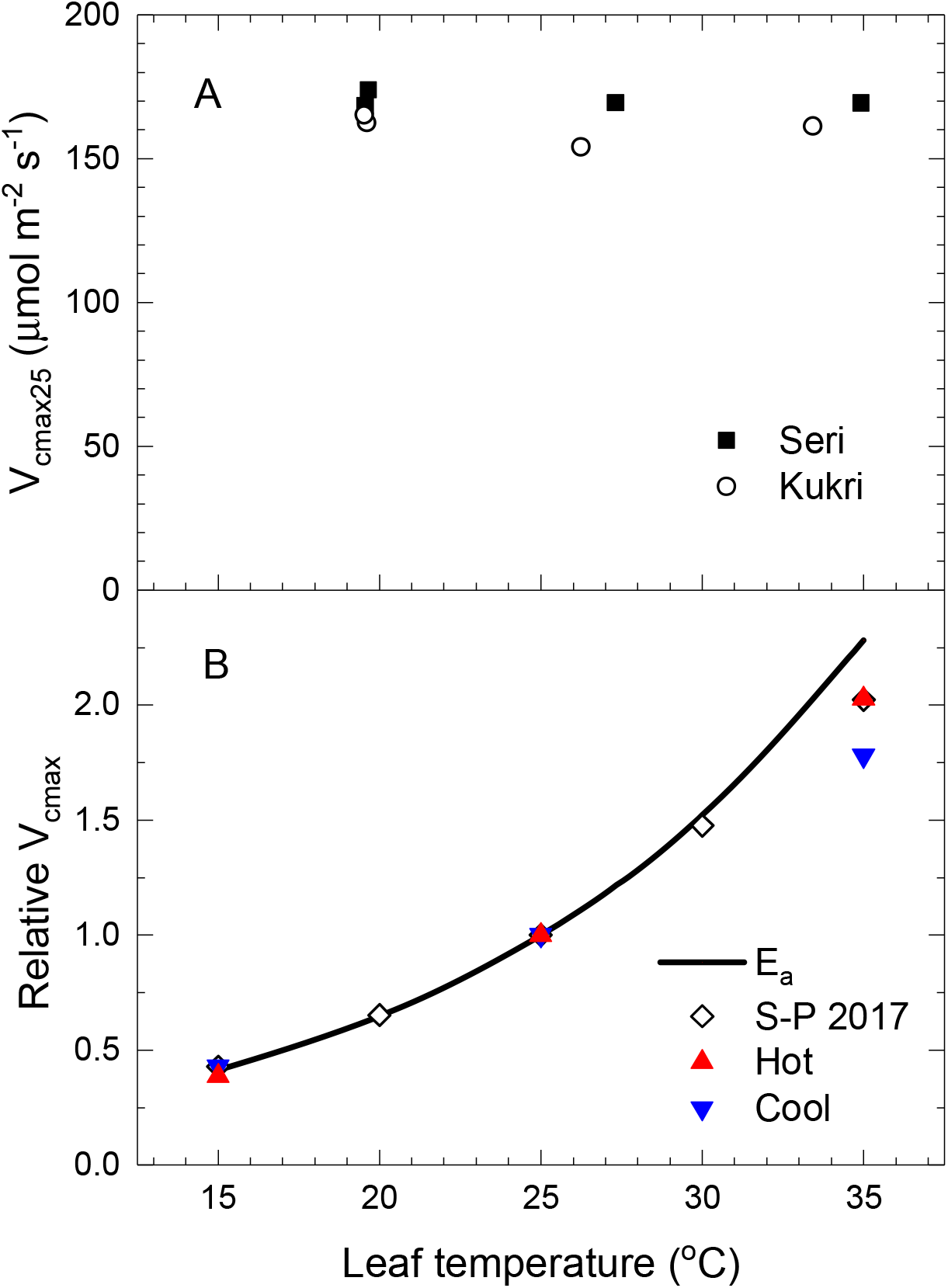
Effects of leaf temperature on Rubisco activity in wheat. A. The maximum rate of carboxylation by Rubisco normalised to 25 °C (*V_cmax25_*) predicted from leaf hyperspectral reflectance measurements at different leaf temperatures. Seven days after anthesis, two wheat genotypes were exposed to a sequence of ambient air temperatures i.e. 15, 25, 35 and 15 °C, in a growth chamber (Expt 1). Symbols represent the mean ± SE of six different leaves from six different plants. B. Temperature response of *V_cmax_* normalised to 1 at 25 °C derived from gas exchange measurements (symbols) or modelled (E_a_ 63 kJ mol^−1^). Data from plants grown under hot (20/15 °C) or cool (32/20 °C) conditions (Expt 2), or from Silva-Perez etal. (2017).

Predicted values for the rate of electron transport, J, were also independent of the leaf temperature when reflectance spectra were measured (Fig. 2A). In this case, the model was built from data collected under a PPFD of 1800 μmol m^−2^ s^−1^ and leaf temperatures mainly at 25°C but ranging up to 32°C. However, in contrast to V_cmax_, J is less sensitive to leaf temperature, increasing by 30% between 15 and 25°C and then plateauing (Fig. 2B). The temperature responses of J were comparable to the previously published data from Silva-Perez et al. (2017). Growth temperature shifted the temperature response. For plants grown under hot conditions, J was less than that from plants grown under cool conditions at 15°C but greater at 35°C. We also measured J using a MultispeQ instrument following the collection of leaf reflectance spectra at each temperature (denoted LET, Fig. 2A). This was measured under a PPFD of 1000 μmol m^−2^ s^−1^, similar to the irradiance in the growth cabinet. As it was collected rapidly, it does not represent the steady state. However, it also indicated that the rate of electron transport was similar between 20 and 30°C, then declined slightly at 35°C.

**Fig. 2.**
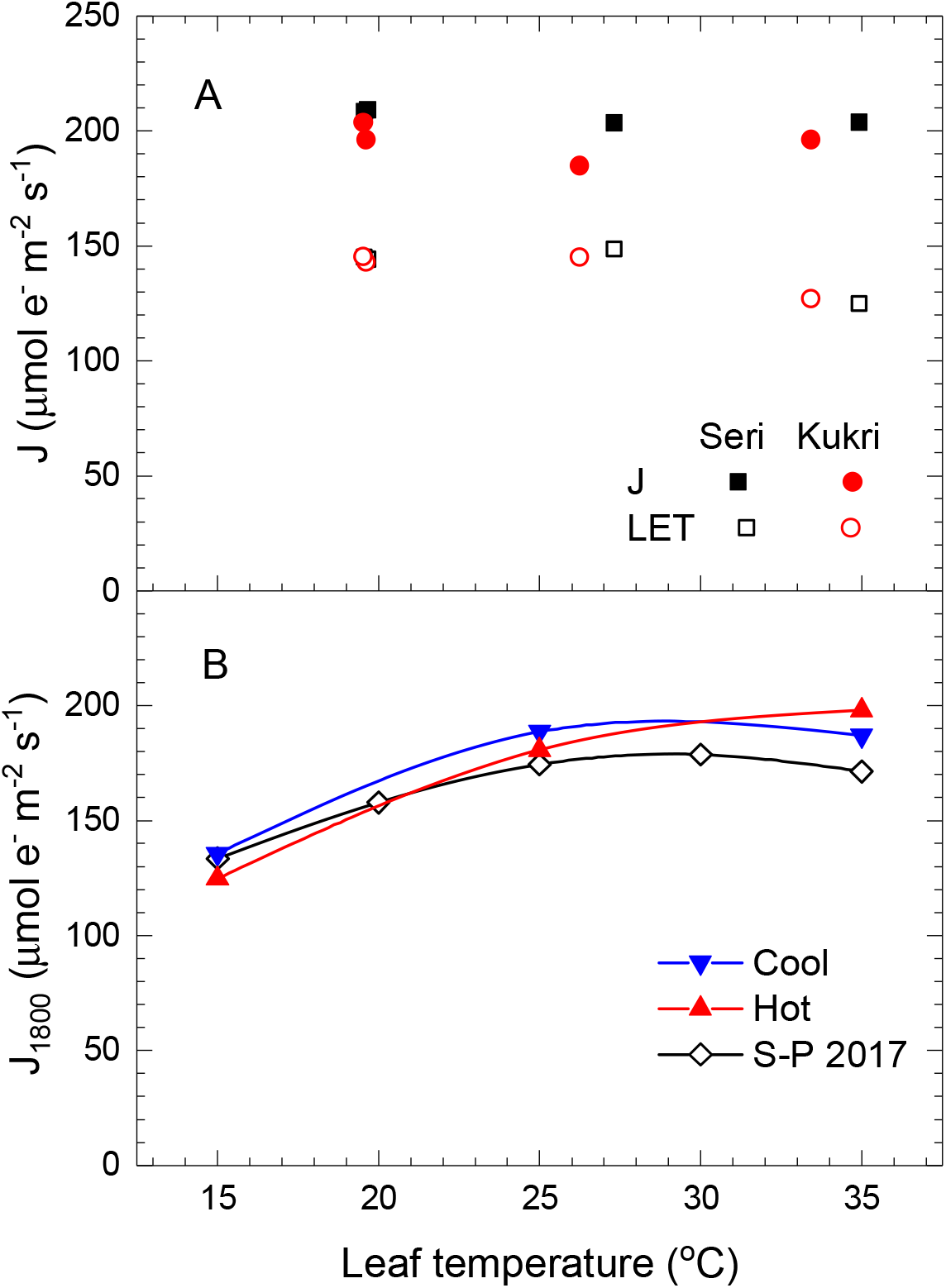
Effects of leaf temperature on electron transport rate in wheat. A. Rate of electron transport (J) predicted from leaf reflectance under a PPFD of 1800 μmol m^−2^ s^−1^ (solid symbols) and linear electron transport (LET) measured using MultispeQ under a PPFD of 1000 μmol m^−2^ s^−1^ (open symbols) at each leaf temperature. Seven days after anthesis, two wheat genotypes were exposed to a sequence of ambient air temperatures i.e. 15, 25, 35 and 15 °C, in a growth chamber (Expt 1). Symbols represent the mean ± SE of six different leaves from six different plants. B. Rates of electron transport calculated from gas exchange measurements made under a PPFD of 1800 μmol m^−2^ s^−1^ from plants grown under cool (20/15 °C) or hot (32/20 °C) conditions (Expt 2), or from Silva Perez et al. (2017).

Predicted values of chlorophyll content were insensitive to the leaf temperature when reflectance spectra were measured (Fig. 3). A similar result was observed for chlorophyll content estimated with the MultispeQ. The absolute values obtained with the MultispeQ were about 20% greater than that predicted from reflectance. The chlorophyll content values are predicted from reflectance using a model built on measurements using SPAD-502 chlorophyll meter (Minolta Camera Co., Ltd, Japan) whereas the MultispeQ uses relative transmissions of red (650 nm) and infrared (940 nm) light. Additionally, MultispeQ has two in built differences from Minolta SPAD; 1) MultispeQ takes a series of transmittance measurements over a range of increasing light intensities, and 2) MultispeQ also averages values over a larger leaf area (~ 1 cm^2^) (Kuhlgert et al. 2016). Nevertheless, additional calibration comparisons were not made as the focus was on temperature. Values predicted for three other leaf traits, leaf dry mass per unit leaf area (LMA), nitrogen content and Rubisco carboxylation capacity normalised to 25°C per unit leaf nitrogen (V_cmax25_/N), were also independent of the leaf temperature when reflectance spectra were measured (Fig. 4). A statistical comparison of the effects of temperature treatments on various measured and predicted leaf traits is also provided separately (Supp. Table 1).

**Fig. 3.**
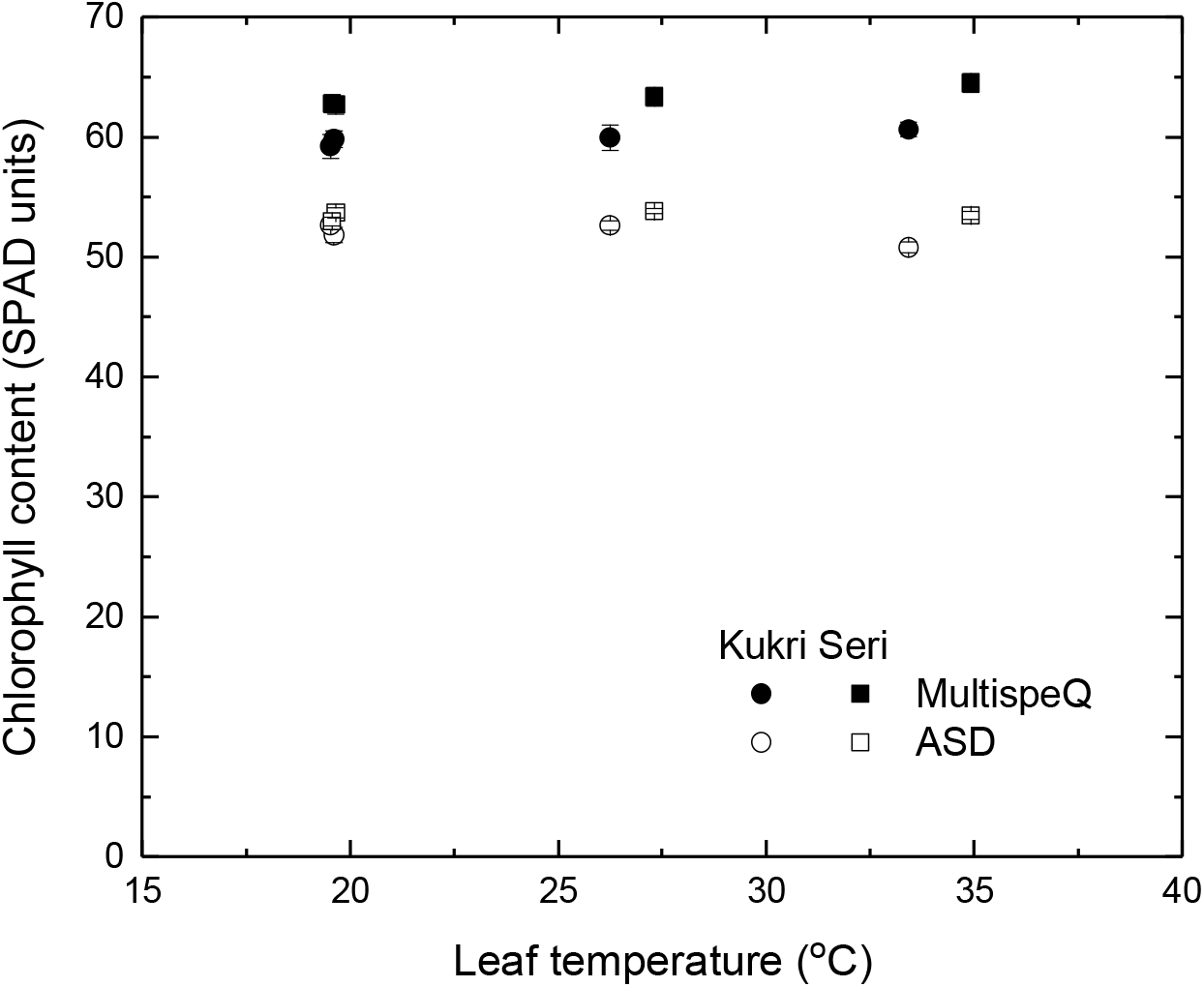
Effects of leaf temperature on estimated chlorophyll content (SPAD units) in wheat. Seven days after anthesis, two wheat genotypes were exposed to a sequence of ambient air temperatures i.e. 15, 25, 35 and 15 °C, in a growth chamber (Expt 1). Symbols represent the mean ± SE of six different leaves from six different plants. Leaf chlorophyll content was estimated using two different devices i.e. direct measurements with MultispeQ or predicted from leaf hyperspectral reflectance measurements made with an ASD FieldSpec Spectroradiometer.

**Fig. 4.**
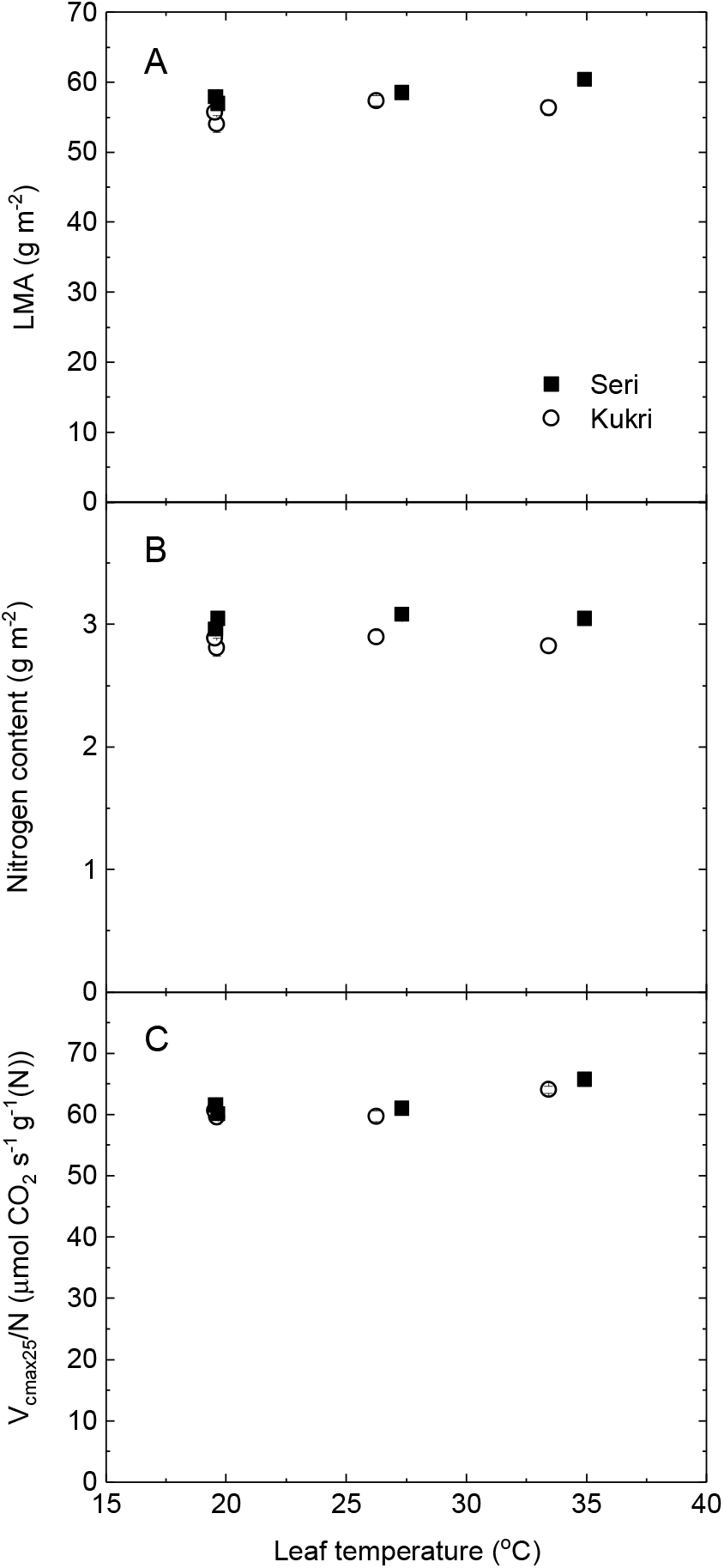
Effects of leaf temperature on parameters predicted from leaf hyperspectral reflectance in wheat. A. Leaf dry mass per unit leaf area (LMA). B. Nitrogen content per unit leaf area. C. Rubisco carboxylation capacity normalised to 25 °C per unit leaf nitrogen (*V_cmax25_*/N). Seven days after anthesis, two wheat genotypes were exposed to a sequence of ambient air temperatures i.e. 15, 25, 35 and 15 °C, in a growth chamber (Expt 1). Symbols represent the mean ± SE of six different leaves from six different plants.

The spectral response of correlations between single wavelength reflectance values and leaf temperature is shown superimposed on the reflectance spectrum (Fig. 5). Clear correlations were observed on the long wavelength shoulders of the two water absorption bands with peaks at 1531 and 2038nm and a third peak in the red edge at 720nm. By sequentially adding weighted reflectance values from single wavelengths, we were able to predict leaf temperature remarkably well with just four wavelengths (Fig. 6).

**Fig. 5.**
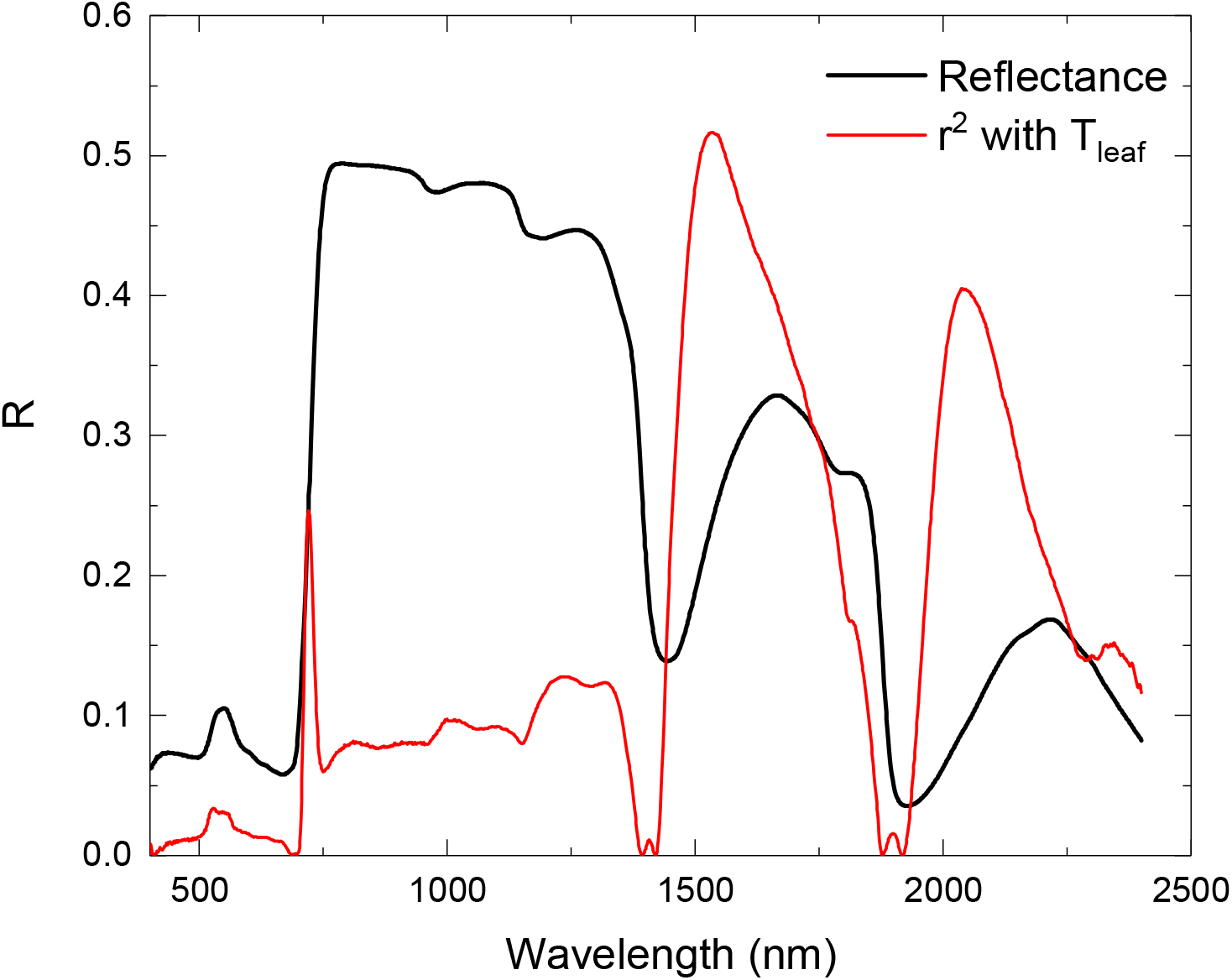
Reflectance spectrum from a wheat leaf measured at 27°C (black thick line) with the spectral response of the correlation coefficient between leaf temperature and leaf reflectance (red thin line) superimposed.

**Fig. 6.**
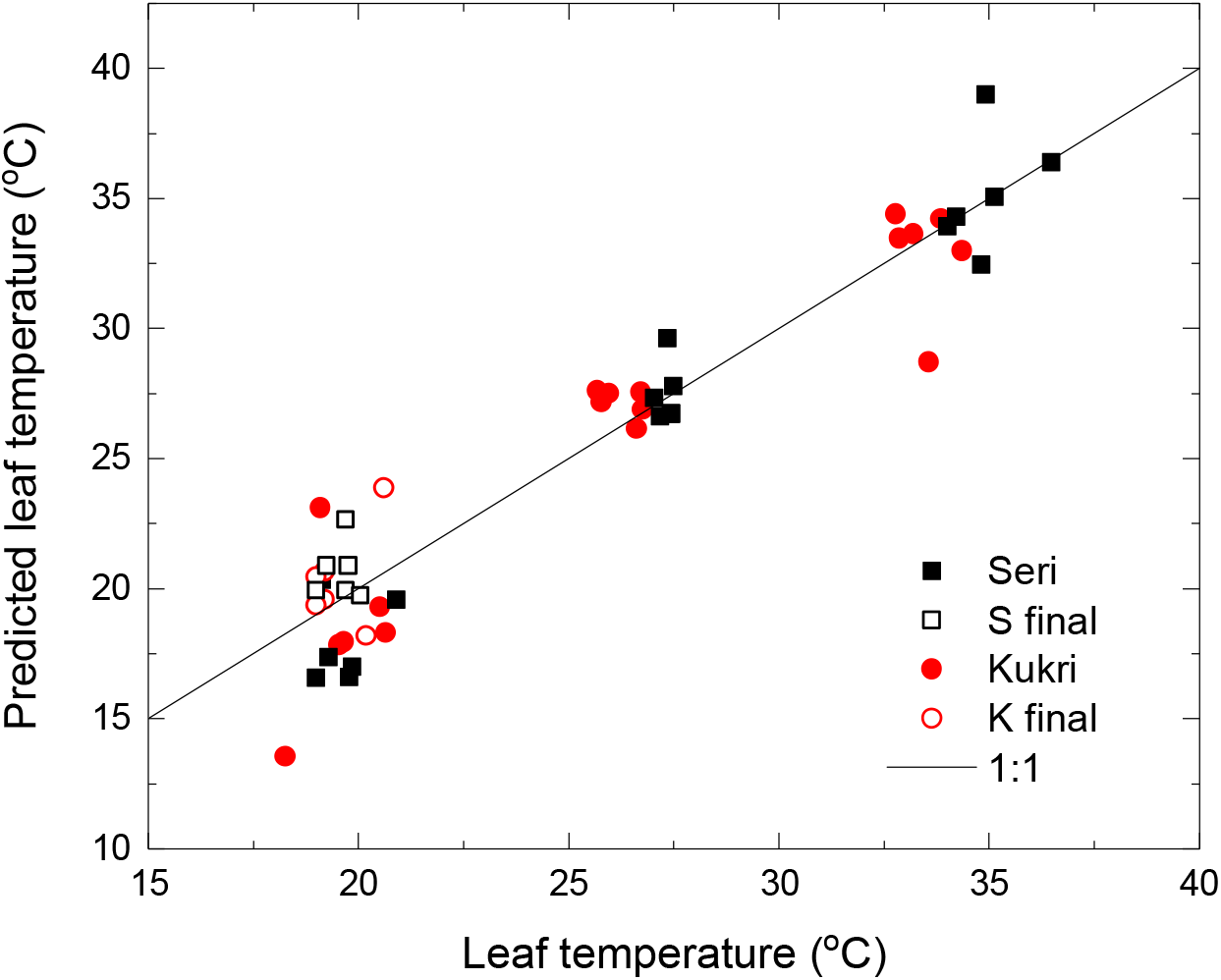
Relationship between leaf temperature predicted from reflectance and measured using MultispeQ in wheat. Two wheat genotypes were exposed to a sequence of ambient air temperatures i.e. 15, 25, 35 and 15 °C which resulted in the leaf temperature increasing from 20 to 35°C (solid symbols) then decreased back to 20°C (open symbols) (Expt 1). Measurements were made on six different leaves from six different plants. The predicted leaf temperature was calculated as: T_leaf_ = 1071.4 R_1531_ – 945.65 R_1400_ – 176.3 R_1507_ + 196.8 R_697_. The fitted regression equation (not shown) was y = 1.015x - 0.4, r^2^ = 0.91.

## Discussion

Leaf hyperspectral reflectance is an optical signal that can provide information remotely and rapidly. With appropriate calibration data obtained from other methods, predictive models can be built for a range of leaf traits. The method has potential for use as a high-throughput tool for phenotyping photosynthetic traits at the leaf and canopy scale. While one might expect leaf reflectance to enable prediction of the amount of a substance e.g. leaf dry mass or nitrogen per unit area, it is harder to understand how physiological processes such as rates of reactions could contribute to reflectance. Thus, with Rubisco being such a major constituent of leaf protein, a reflectance model could arise from a signal associated with leaf protein or nitrogen, as argued by Dechant et al. (2017). If this was the case, then changing leaf temperature would alter Rubisco activity but not Rubisco content. However, Serbin et al. (2012) successfully included variation in leaf temperature to derive a model predicting Rubisco activity, demonstrating that the model was not fundamentally associated with another constituent such as nitrogen. In contrast to gas exchange measurements, where leaf temperature is measured directly to enable the calculation of stomatal conductance, leaf temperature is not generally measured directly during the collection of hyperspectral reflectance. Consequently, it is necessary to consider how leaf temperature affects the estimation of physiological traits, such as V_cmax_ and J, using leaf hyperspectral reflectance.

### Parameters independent of temperature

A leaf structural property that has been widely reported is leaf dry mass per unit leaf area as it is easy to measure and relates to lifespan and other traits (Wright *et al*., 2004). Robust predictions of LMA can be made from hyperspectral reflectance data (Ecarnot *et al*., 2013; Serbin *et al*., 2012; Silva-Perez *et al*., 2018). As LMA is a leaf property that would not change in response to short term changes in temperature, models predicting LMA from hyperspectral reflectance should also be insensitive to the temperature of the leaf during measurement. This was found to be true for LMA (Fig. 4A) as well as for other leaf constituents, chlorophyll (Fig. 3) and nitrogen (Fig. 4B).

Photosynthesis is a process involving many constituents, but has been successfully modelled in C_3_ plants by considering the properties of Rubisco (Farquhar *et al*., 1980). Knowing the amount of Rubisco per unit leaf area, its properties and a few assumptions, it is possible to predict photosynthetic responses to irradiance, atmospheric CO_2_ and temperature. The amount of Rubisco is unlikely to vary significantly during short term changes in leaf temperature, but the carboxylase activity is strongly temperature dependent (Badger and Collatz, 1977; Bernacchi *et al*., 2001; Sharwood *et al*., 2016). Therefore, models using reflectance to predict Rubisco content or activity normalised to a fixed temperature should be independent of leaf temperature. Indeed, this was what we observed (Fig. 1A).

### Parameters that vary with temperature

Enzyme activities vary with temperature which can be described with the Arrhenius equation through the energy of activation term E_a_. V_cmax_ is the product of Rubisco content and catalytic rate. It seems possible that models predicting V_cmax_ from reflectance (Ainsworth *et al*., 2014; Serbin *et al*., 2012) may contain two components, one that is independent of temperature (representing Rubisco protein content) and another that varies with leaf temperature. Immediately prior to placing the leaf into the clip for measuring leaf reflectance, Ainsworth et al. (2014) measured leaf temperature with an infrared thermometer, but they only report the relationship between predicted V_cmax_ and leaf temperature. We, therefore, looked at our reflectance spectra collected at different leaf temperatures to see if we could predict leaf temperature. Predicted leaf temperature, using reflectance of just four wavelengths, clearly correlated with measured leaf temperature, with an r^2^ of 0.91 (Fig. 6). It is unlikely that this equation has general application as it arose from only two genotypes measured under one environment and it is known that the power and generality of models predicting dark respiration (Coast *et al*., 2019) and Vcma×25 (Wu *et al*., 2019) improved as the diversity of calibration data increased. However, the point is that leaf temperature apparently can be extracted from reflectance spectra which could explain how models can predict reaction rates from reflectance.

The Arrhenius equation predicts an exponential increase in rate with increasing temperature, whereas models calculating parameters from reflectance sum linear weightings of each reflectance at each wavelength and would, therefore, have linear responses to temperature. The difference between an exponential and a linear relationship may not be very noticeable over a narrow temperature range. In the case of V_cma_χ, values derived from gas exchange deviated below the Arrhenius function at 35°C (Fig. 1B), such that a linear function would fit the data well between 15 and 35°C. There are also indications that a single value for E_a_ may not be appropriate across the temperature range from 10 to 40°C. Sharwood et al. (2016) found it necessary to use lower values for E_a_ at leaf temperatures above 25°C. The rate of electron transport, J, also varies with temperature (Bernacchi *et al*., 2003; June *et al*., 2004; Medlyn *et al*., 2002) but reaches a maximum around 30°C before decreasing again. As a result, the change in J between 20 and 35°C is less pronounced than for V_cmax_. It is therefore uncertain whether models predicting J from reflectance (Dechant *et al*., 2017; Serbin *et al*., 2012; Silva-Perez *et al*., 2018) would contain components that vary with temperature. In the case of J for wheat (Silva-Perez *et al*., 2018), the reflectance model was built with data that varied little in leaf temperature and the predicted values of J were found to be unaffected by leaf temperature when reflectance was measured (Fig. 2A). However, given that J did not vary greatly over this temperature range (Fig. 2), this may not be a very rigorous test. By contrast, as Serbin et al. (2012) deliberately used variation in leaf temperature to generate a broader spread in J to build their reflectance model – testing their function with multiple spectra obtained from a leaf measured over a range of temperatures could be informative.

## Conclusion

Leaf temperature varying between 20 to 35°C during the measurement of leaf reflectance did not affect predicted values of leaf traits (V_cmax25_, chlorophyll and nitrogen contents per unit area, LMA and V_cmax_/N), for wheat. It was possible to extract leaf temperature from reflectance spectra which may explain how models that predict rates that vary with temperature (V_cmax_, J, dark respiration) could arise. Models predicting traits that vary with leaf temperature should be tested using multiple measurements from each leaf covering a range of temperature. Reflectance appears to have the potential to predict leaf temperature, but to construct a robust model would require calibration with a broader set of experiments.

## Acknowledgements

This work was made possible by funding from the GRDC through grant ANU00025, which is part of IWYP60 Using next generation approaches to exploit phenotypic variation in photosynthetic efficiency to increase wheat yield, financial support from the Australian Government through the Australian Research Council Centre of Excellence for Translational Photosynthesis (CE140100015), and a Cooperative Laboratory Study Program grant to YN.

## Supplementary material

**Supplementary. Table 1.** Effects of leaf temperature on leaf physiological traits measured using MulitispeQ (leaf temperature, LET, relative chlorophyll) or predicted from leaf hyperspectral reflectance (V_cmax25_, J, chlorophyll content, LMA, nitrogen content, V_cmax25_/N) in leaves of two wheat genotypes exposed to a sequence of ambient air temperatures i.e. 15, 25, 35 and 15 °C, in a growth chamber (Expt 1). Symbols represent the mean ± SE of six different leaves from six different plants. Sequential measurements were made on each leaf, seven days after anthesis.

| Genotype | Ambient Air Temperature | Measured traits using MultispeQ |  |  | Predicted traits using leaf hyperspectral reflectance |  |  |  |  |  |
| --- | --- | --- | --- | --- | --- | --- | --- | --- | --- | --- |
| | | Leaf Temperature (°C) | LET ($\mu\text{mol e}^- \text{m}^{-2} \text{s}^{-1}$) | Relative Chlorophyll (SPAD units) | $V_{\text{cmax}25}$ ($\mu\text{mol CO}_2 \text{s}^{-1} \text{g}^{-1} (\text{N})$) | J ($\mu\text{mol e}^- \text{m}^{-2} \text{s}^{-1}$) | Chlorophyll content (SPAD units) | LMA ($\text{g m}^{-2}$) | Nitrogen content ($\text{g m}^{-2}$) | $V_{\text{cmax}25}/N$ ($\mu\text{mol CO}_2 \text{s}^{-1} \text{g}^{-1} (\text{N})$) |
| Kukri | 15 °C | 19.6 ± 0.4 | 142.9 ± 7.0 | 59.8 ± 0.7 | 162.5<sup>a</sup> ± 1.7 | 196.2<sup>ab</sup> ± 3.6 | 51.8<sup>ab</sup> ± 0.6 | 54.0 ± 1.2 | 2.81 ± 0.07 | 59.6<sup>b</sup> ± 1.2 |
| | 25 °C | 26.2 ± 0.2 | 145.2 ± 9.2 | 59.9 ± 1.0 | 154.2<sup>b</sup> ± 1.8 | 184.9<sup>b</sup> ± 3.7 | 52.6<sup>a</sup> ± 0.4 | 57.4 ± 0.8 | 2.90 ± 0.05 | 59.7<sup>b</sup> ± 0.9 |
| | 35 °C | 33.4 ± 0.3 | 127.0 ± 4.3 | 60.6 ± 0.6 | 161.3<sup>a</sup> ± 2.2 | 196.3<sup>ab</sup> ± 4.5 | 50.8<sup>b</sup> ± 0.4 | 56.3 ± 1.1 | 2.82 ± 0.06 | 64.1<sup>a</sup> ± 0.6 |
| | 15 °C | 19.5 ± 0.3 | 145.4 ± 8.1 | 59.2 ± 1.0 | 165.2<sup>a</sup> ± 2.6 | 203.8<sup>a</sup> ± 5.0 | 52.7<sup>a</sup> ± 0.3 | 55.7 ± 1.1 | 2.89 ± 0.05 | 60.6<sup>b</sup> ± 1.0 |
|  | LSD (5%) |  | n.s. | n.s. | 6.2** | 12.5* | 1.3* | n.s. | n.s. | 2.8* |
| Seri | 15 °C | 19.7 ± 0.3 | 144.2<sup>a</sup> ± 5.2 | 62.7 ± 0.8 | 173.9 ± 2.4 | 209.2 ± 4.0 | 53.7 ± 0.4 | 56.9<sup>b</sup> ± 0.7 | 3.05 ± 0.05 | 60.1<sup>b</sup> ± 0.8 |
| | 25 °C | 27.3 ± 0.1 | 148.6<sup>a</sup> ± 6.1 | 63.3 ± 0.8 | 169.5 ± 2.2 | 203.5 ± 3.7 | 53.8 ± 0.2 | 58.5<sup>ab</sup> ± 0.6 | 3.08 ± 0.05 | 61.0<sup>b</sup> ± 0.7 |
| | 35 °C | 34.9 ± 0.4 | 124.9<sup>b</sup> ± 3.2 | 64.5 ± 0.8 | 169.4 ± 2.2 | 203.7 ± 4.3 | 53.5 ± 0.4 | 60.4<sup>a</sup> ± 0.7 | 3.05 ± 0.05 | 65.7<sup>a</sup> ± 0.7 |
| | 15 °C | 19.6 ± 0.2 | 145.1<sup>a</sup> ± 6.5 | 62.8 ± 0.7 | 168.4 ± 2.2 | 208.5 ± 3.7 | 53.0 ± 0.3 | 57.9<sup>b</sup> ± 0.9 | 2.96 ± 0.04 | 61.5<sup>b</sup> ± 0.5 |
|  | LSD (5%) |  | 15.9* | n.s. | n.s. | n.s. | n.s. | 2.1* | n.s. | 2.04*** |

\*\*\* = $P < 0.001$ , \*\* = $P < 0.01$ , \* = $P < 0.05$ , n.s. = non-significant

## References

Ainsworth EA, Serbin SP, Skoneczka JA, Townsend PA. 2014. Using leaf optical properties to detect ozone effects on foliar biochemistry. PHOTOSYNTHESIS RESEARCH 119, 65–76.

Badger M, Collatz GJ. 1977. Studies on the kinetic mechanism of ribulose-1,5-bisphosphate carboxylase and oxygenase reactions, with particular reference to the effect of temperature on kinetic parameters. Carnegie Yearbook, Vol. 76, 355–361.

Beche E, Benin G, da Silva CL, Munaro LB, Marchese JA. 2014. Genetic gain in yield and changes associated with physiological traits in Brazilian wheat during the 20th century. European Journal of Agronomy 61, 49–59.

Bernacchi CJ, Pimentel C, Long SP. 2003. In vivo temperature response functions of parameters required to model RuBP-limited photosynthesis. Plant, Cell & Environment 26, 1419–1430.

Bernacchi CJ, Singsaas EL, Pimentel C, Portis AR, Long SP. 2001. Improved temperature response functions for models of Rubisco-limited photosynthesis. Plant, Cell & Environment 24, 253–259.

Coast O, Shah S, Ivakov A, Gaju O, Wilson PB, Posch BC, Bryant CJ, Negrini ACA, Evans JR, Condon AG, Silva-Pérez V, Reynolds MP, Pogson BJ, Millar AH, Furbank RT, Atkin OK. 2019. Predicting dark respiration rates of wheat leaves from hyperspectral reflectance. Plant, Cell & Environment 42, 2133–2150.

Dechant B, Cuntz M, Vohland M, Schulz E, Doktor D. 2017. Estimation of photosynthesis traits from leaf reflectance spectra: Correlation to nitrogen content as the dominant mechanism. Remote Sensing of Environment 196, 279–292.

Driever SM, Lawson T, Andralojc PJ, Raines CA, Parry MAJ. 2014. Natural variation in photosynthetic capacity, growth, and yield in 64 field-grown wheat genotypes. JOURNAL OF EXPERIMENTAL BOTANY.

Ecarnot M, Compan F, Roumet P. 2013. Assessing leaf nitrogen content and leaf mass per unit area of wheat in the field throughout plant cycle with a portable spectrometer. Field Crops Research 140, 44–50.

Evans JR, von Caemmerer S. 2013. Temperature response of carbon isotope discrimination and mesophyll conductance in tobacco. Plant Cell & Environment 36, 745–756.

Farquhar GD, von Caemmerer S, Berry JA. 1980. A biochemical model of photosynthetic CO_2_ assimilation in leaves of C_3_ species. PLANTA 149, 78–90.

Fischer RA, Rees D, Sayre KD, Lu ZM, Condon AG, Saavedra AL. 1998. Wheat yield progress associated with higher stomatal conductance and photosynthetic rate, and cooler canopies. Crop Science 38, 1467–1475.

Gaju O, DeSilva J, Carvalho P, Hawkesford MJ, Griffiths S, Greenland A, Foulkes MJ. 2016. Leaf photosynthesis and associations with grain yield, biomass and nitrogen-use efficiency in landraces, synthetic-derived lines and cultivars in wheat. Field Crops Research 193, 1–15.

Heckmann D, Schluter U, Weber APM. 2017. Machine learning techniques for predicting crop photosynthetic capacity from leaf reflectance spectra. Molecular Plant 10, 878–890.

June T, Evans JR, Farquhar GD. 2004. A simple new equation for the reversible temperature dependence of photosynthetic electron transport: a study on soybean leaf. FUNCTIONAL PLANT BIOLOGY 31, 275–283.

Kuhlgert S, Austic G, Zegarac R, Osei-Bonsu I, Hoh D, Chilvers MI, Roth MG, Bi K, TerAvest D, Weebadde P, Kramer DM. 2016. MultispeQ Beta: a tool for large-scale plant phenotyping connected to the open PhotosynQ network. Royal Society Open Science 3.

Meacham-Hensold K, Montes CM, Wu J, Guan KY, Fu P, Ainsworth EA, Pederson T, Moore CE, Brown KL, Raines C, Bernacchi CJ. 2019. High-throughput field phenotyping using hyperspectral reflectance and partial least squares regression (PLSR) reveals genetic modifications to photosynthetic capacity. Remote Sensing of Environment 231.

Medlyn BE, Dreyer E, Ellsworth D, Forstreuter M, Harley PC, Kirschbaum MUF, Le Roux X, Montpied P, Strassemeyer J, Walcroft A, Wang K, Loustau D. 2002. Temperature response of parameters of a biochemically based model of photosynthesis. II. A review of experimental data. Plant, Cell & Environment 25, 1167–1179.

Parry MAJ, Reynolds M, Salvucci ME, Raines C, Andralojc PJ, Zhu ×-G, Price GD, Condon AG, Furbank RT. 2011. Raising yield potential of wheat. II. Increasing photosynthetic capacity and efficiency. JOURNAL OF EXPERIMENTAL BOTANY 62, 453–467.

R. 2013. R: A language and environment for statistical computing. R Foundation for Statistical Computing, Vienna, Austria.

Reynolds M, Foulkes MJ, Slafer GA, Berry P, Parry MAJ, Snape JW, Angus WJ. 2009. Raising yield potential in wheat. JOURNAL OF EXPERIMENTAL BOTANY 60, 1899–1918.

Sadras VO, Lawson C, Montoro A. 2012. Photosynthetic traits in Australian wheat varieties released between 1958 and 2007. Field Crops Research 134, 19–29.

Serbin SP, Dillaway DN, Kruger EL, Townsend PA. 2012. Leaf optical properties reflect variation in photosynthetic metabolism and its sensitivity to temperature. JOURNAL OF EXPERIMENTAL BOTANY 63, 489–502.

Sharwood RE, Ghannoum O, Kapralov MV, Gunn LH, Whitney SM. 2016. Temperature responses of Rubisco from Paniceae grasses provide opportunities for improving C3 photosynthesis. Nature Plants 2, 16186.

Shearman VJ, Sylvester-Bradley R, Scott RK, Foulkes MJ. 2005. Physiological processes associated with wheat yield progress in the UK. Crop Science 45, 175–185.

Silva-Pérez V, De Faveri J, Molero G, Deery DM, Condon AG, Reynolds MP, Evans JR, Furbank RT. 2019. Genetic variation for photosynthetic capacity and efficiency in spring wheat. JOURNAL OF EXPERIMENTAL BOTANY 71, 2299–2311.

Silva-Perez V, Furbank RT, Condon AG, Evans JR. 2017. Biochemical model of C-3 photosynthesis applied to wheat at different temperatures. PLANT CELL AND ENVIRONMENT 40, 1552–1564.

Silva-Perez V, Molero G, Serbin SP, Condon AG, Reynolds MP, Furbank RT, Evans JR. 2018. Hyperspectral reflectance as a tool to measure biochemical and physiological traits in wheat. JOURNAL OF EXPERIMENTAL BOTANY 69, 483–496.

Wright IJ, Reich PB, Westoby M, Ackerly DD, Baruch Z, Bongers F, Cavender-Bares J, Chapin T, Cornelissen JHC, Diemer M, Flexas J, Garnier E, Groom PK, Gulias J, Hikosaka K, Lamont BB, Lee T, Lee W, Lusk C, Midgley J J, Navas M-L, Niinemets U, Oleksyn J, Osada N, Poorter H, Poot P, Prior L, Pyankov VI, Roumet C, Thomas SC, Tjoelker MG, Veneklaas EJ, Villar R. 2004. The worldwide leaf economics spectrum. Nature 428, 821–827.

Wu J, Rogers A, Albert LP, Ely K, Prohaska N, Wolfe BT, Oliveira RC, Saleska SR, Serbin SP. 2019. Leaf reflectance spectroscopy captures variation in carboxylation capacity across species, canopy environment and leaf age in lowland moist tropical forests. NEW PHYTOLOGIST 224, 663–674.

Yao YR, Lv LH, Zhang LH, Yao HP, Dong ZQ, Zhang JT, Ji JJ, Jia XL, Wang HJ. 2019. Genetic gains in grain yield and physiological traits of winter wheat in Hebei Province of China, from 1964 to 2007. Field Crops Research 239, 114–123.

Yendrek CR, Tomaz T, Montes CM, Cao Y, Morse AM, Brown PJ, McIntyre LM, Leakey ADB, Ainsworth EA. 2017. High-throughput phenotyping of maize leaf physiological and biochemical traits using hyperspectral reflectance. PLANT PHYSIOLOGY 173, 614–626.

